# Optimized Repli-seq: Improved DNA Replication Timing Analysis by Next-Generation Sequencing

**DOI:** 10.1101/2022.04.26.489601

**Authors:** Juan Carlos Rivera-Mulia, Claudia Trevilla-Garcia, Santiago Martinez-Cifuentes

## Abstract

The human genome is divided into functional units that replicate at specific times during S-phase. This temporal program is known as replication timing (RT) and is coordinated with the spatial organization of the genome and transcriptional activity. RT is also cell type-specific, dynamically regulated during development, and alterations in RT are observed in multiple diseases. Thus, precise measure of RT is critical to understand the role of RT in gene function regulation. Distinct methods for assaying the RT program exist; however, conventional methods require thousands of cells as input, prohibiting its applicability to samples with limited cell numbers such as those from disease patients or from early developing embryos. Although single-cell analysis of RT has been developed as an alternative, these methods are low throughput and produce low resolution data. Here, we developed an improved method to measure RT genome-wide that enables high resolution analysis of low input samples. This method incorporates direct cell sorting into lysis buffer, as well as DNA fragmentation and library preparation in a single tube, resulting in higher yields, increased quality, and reproducibility with decreased costs. We also performed a systematic data processing analysis to provide standardized parameters for RT measurement. This optimized method facilitates RT analysis and will enable its application to a broad range of studies investigating the role of RT in gene expression, nuclear architecture, and disease.

## Introduction

The eukaryotic genome is partitioned into submegabase units that segregate to distinct nuclear compartments and replicate in a specific order during S-phase. This replication timing (RT) program is coordinated with the transcriptional activity, with early replication corresponding to active chromatin and repressive chromatin regions replicating late in S-phase (Rivera-Mulia and Gilbert 2016a, b). RT also correlates with the 3D genome organization and replication domains closely align with the self-interacting topologically associating domains (TADs) identified by chromosome conformation capture technologies (Ryba et al. 2010; Yaffe et al. 2010; Dixon et al. 2012; Moindrot et al. 2012; Pope et al. 2014; Miura et al. 2019). RT is cell type-specific and is dynamically remodeled during development (Hiratani et al. 2008, 2010; Rivera-Mulia et al. 2015, 2019a). Moreover, alterations in RT are associated with many diseases (Ryba et al. 2012; Sasaki et al. 2017; Rivera-Mulia et al. 2017, 2018b, 2019b; Zhang et al. 2020), highlighting the importance of understanding the role of RT in gene regulation.

Distinct methods to measure RT genome-wide have been developed allowing the genome-wide assessment of the temporal replication program. Recently, we developed Repli-seq (Marchal et al. 2018), for RT analysis by next-generation sequencing (NGS) as an adaptation of the previous methods based on microarray hybridization (Schübeler et al. 2002; MacAlpine et al. 2004; Farkash-Amar et al. 2008; Hiratani et al. 2008, 2010; Ryba et al. 2011). In Repli-seq, cells are pulse-labeled with nucleotide analogs to label nascent DNA. Subsequently, populations of early and late S-phase cells are isolated by flow cytometry and replicated DNA is purified and sequenced (Hansen et al. 2010; Marchal et al. 2018). This method is very robust and highly reproducible, but a limitation is that it requires a large number of proliferating cells (Marchal et al. 2018; Zhao et al. 2020). In fact, processing of rare, limited samples, such as those derived from disease patients, is challenging and usually, high quality data can be obtained only from a small subset of samples (Sasaki et al. 2017; Rivera-Mulia et al. 2017, 2019b). Alternative approaches have been developed, such as measuring RT as the ratio of sequenced reads from S-phase versus G1 cells, which relies on the increasing DNA copy number during S-phase (Woodfine et al. 2004; Desprat et al. 2009; Koren et al. 2014). A disadvantage of this method is that, since the maximum copy number difference is two, the resulting dynamic range (two-fold) is much lower than that in E/L Repli-seq (4,000-fold). A third method for ensemble analysis of RT, is the calculation of copy number variation from deep whole-genome sequencing (Koren et al. 2014, 2021). This alternative allows measuring RT without the need of cell purification by flow cytometry; however, it requires costly deep sequencing (≥30X coverage) and the dynamic range of the data obtained is even lower than that of the S/G1 Repli-seq. Single-cell Repli-seq (scRepli-seq) has also been achieved (Dileep and Gilbert 2018; Takahashi et al. 2019; Miura et al. 2020; Bartlett et al. 2022), and it allows estimation of cell-to-cell variability in RT. However, scRepli-seq is low throughput, very laborious and expensive, requires the generation of multiple single-cell libraries per sample, and has low resolution. Here, we developed an optimized Repli-seq method that allows measurement of RT from low input samples without compromising resolution.

## Results and discussion

### Repli-seq protocol overview

Standard Repli-seq requires incubating cell samples for a short period (~2 hours) with a nucleotide analogue (most commonly 5-Bromodeoxyuridine-BrdU) to label the replicating DNA. Then, cells are fixed and cell populations in early and late S-phase are purified by flow cytometry based on the DNA content. DNA content is measured based on fluorescence intensity of DNA-binding dyes; the most commonly used is propidium iodide (PI). Since PI also binds RNA, sample preparation for FACS of PI-stained cells also requires treatment with RNase. Sorted cells in early and late S-phase fraction are then lysed and genomic DNA is isolated. Library preparation for NGS is performed and nascent DNA is purified by immunoprecipitation using antibodies against BrdU. Libraries are then amplified and after sequencing, mapped reads are used to calculate the RT as the Log2 ratio between early and late fractions (**Fig. 1**).

**Fig. 1.**
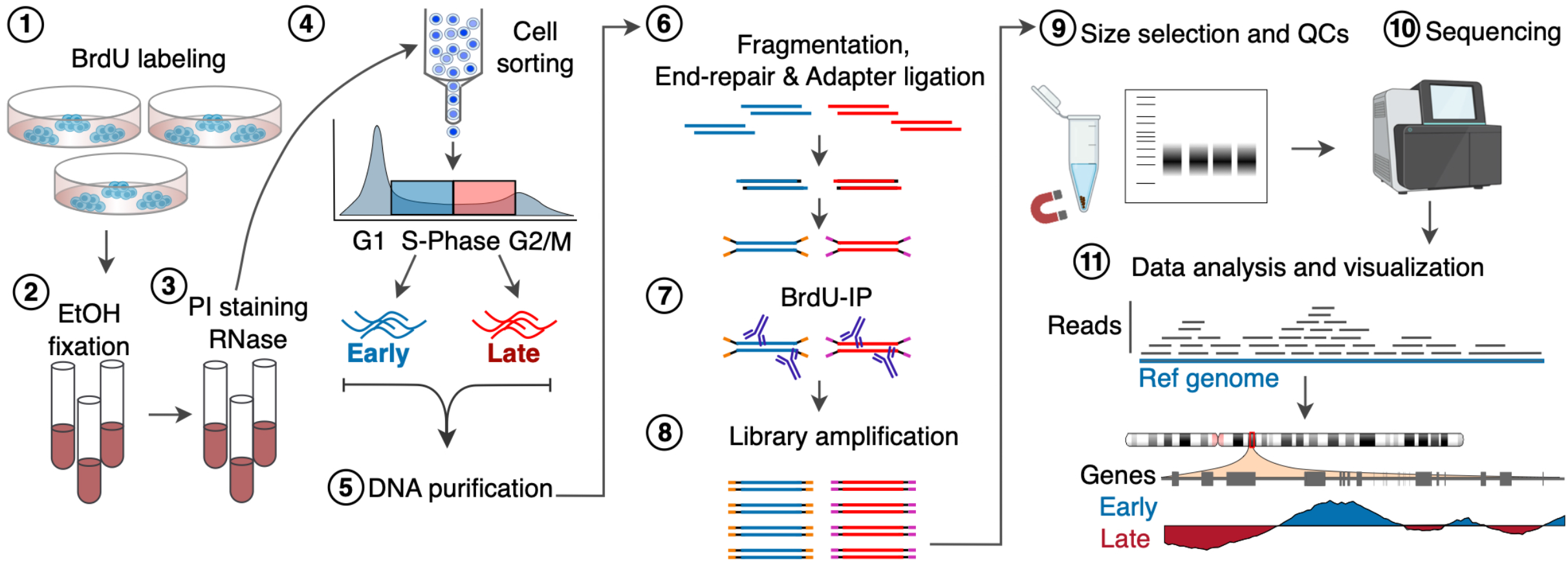
Overview of the optimized Repli-seq method. BrdU-labeled cells (Step 1) are dissociated, fixed in ethanol (Step 2), stained with PI and treated with RNase (Step 3). Early and late S-phase fraction cells are obtained by flow cytometry (Step 4). DNA from FACS sorted cells is purified (Step 5) and used for library preparation for next generation sequencing (Step 6). Libraries are then immunoprecipitated with anti-BrdU antibody (Step 7) and amplified (Step 8). After quality controls (Step 9) and sequencing (Step 10), RT profiles are calculated by the Log2 ratio between mapped reads of early/late fractions (Step 11).

### An improved method for genomic DNA purification from proliferating cells

Previously, we developed the standard Repli-seq method and variations of this method that can help analyze more challenging samples, such as those limited samples obtained from leukemia patients (Marchal et al. 2018; Rivera-Mulia et al. 2019b). In this method, samples are sorted into large aliquots of ~120,000 cells of each early and late S-phase fractions. Then, sorted cells are lysed and digested with SDS and proteinase K for 2 hours, pelleted and divided into aliquots of 20,000 to 80,000 cells (Ryba et al. 2011; Marchal et al. 2018; Rivera-Mulia et al. 2019b; Zhao et al. 2020). Finally, genomic DNA is purified using silica-based nucleic acid methods. However, this workflow is time consuming, requires thousands of purified proliferating cells and multiple tube-to-tube sample transfers that cause a significant sample loss (**Fig. 2A**). In fact, nucleic acid purification from sorted cells using this standard method consistently yields <25% of the total expected DNA (considering 6pg of DNA per diploid mammalian cell). To avoid sample loss, we optimized the sample processing by: 1) sorting cells directly into the lysis buffer; and 2) reducing multiple sample transfers by excluding the digestion step with proteinase K. To achieve this, we collected equal cell numbers of early and late fractions by sorting samples directly into the lysis buffer (**Fig. 2A**). Although any column-based DNA purification method should work, we optimized our protocol using the Zymo Quick-DNA purification, as it allows sample elution in volumes as low as 10 ul, which are directly compatible with library preparation without requiring further sample concentration. In addition, the genomic lysis buffer from this kit allows cell lysis without the need for proteinase K digestion. To validate our optimized protocol, we processed two human cell lines (H9 embryonic stem cells and U2OS osteosarcoma cells), as well as mouse embryonic fibroblasts (MEFs) by the standard and the optimized protocols. H9 and U2OS cell lines were chosen to analyze human samples with distinct levels of proliferating cells (H9 typically contain ≥40 % of cells in S-phase, while U2OS cells have a S-phase population ≤15 %), and MEFs cells were exploited to analyze our method in distinct organism. Cell cycle profiles and sorting gates are shown in **Fig. 2B**. Then, we sorted 20,000 cells (minimum number of cells recommended in the standard Repli-seq) of each early and late S-phase fractions, either into 400ul of PBS/FBS buffer or directly into the genomic lysis buffer. Cells sorted into PBS/FBS were processed by the standard method, i.e., sorted, digested by 2 hours with proteinase K, pelleted and then the DNA was purified. Cells sorted directly into the lysis buffer were processed immediately for genomic DNA purification following the manufacturer’s instructions. Increased yields resulted by using our optimized protocol, and 3-fold increase in DNA yield was consistently achieved for all cell lines processed (**Fig. 2C**). Moreover, our optimized protocol reduced the DNA preparation from FACS-sorted samples for Repli-seq from ≥ 3 hours to less than 20 minutes. Thus, the modifications incorporated into the optimized protocol allowed improved yields and a rapid, single-tube isolation of high-quality DNA for genome-wide analysis of RT.

**Fig. 2.**
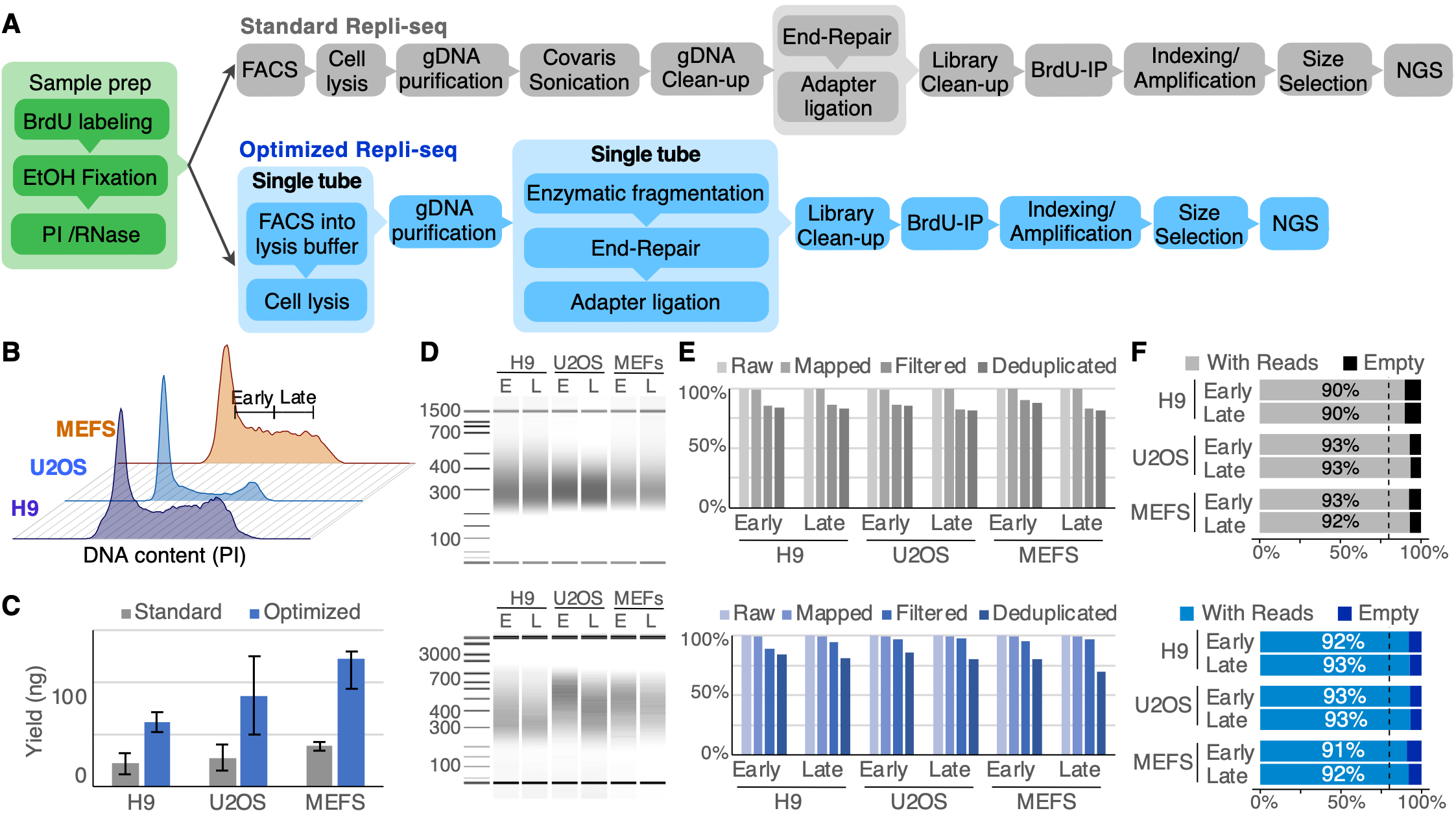
Comparison of standard and optimized Repli-seq methods. A). Workflow of standard and optimized Repli-seq. Direct sorting into the lysis buffer, as well as performing fragmentation and library preparation in the same tube allows faster sample processing with increased yields of input DNA. B). Cell cycle analysis of the distinct cell lines analyzed. Cells were stained with PI, treated with RNase and cell cycle was analyzed based on DNA content by flow cytometry. Gates used to purify cells in early and late S-phase are shown. C) Increased genomic DNA yields (3-fold) were observed using the optimized Repli-seq method for all cell lines tested. Bars represent the average of 4 replicates and standard deviation is shown. D) Comparison of size distribution after genomic DNA fragmentation using standard (top) and optimized (bottom) Repli-seq methods. E) Comparison of NGS reads obtained by the standard (gray bars) and optimized (blue bars) Repli-seq methods. All samples were sequenced at ~8M reads per library using pair-end sequencing at 50 bp in the Illumina NovaSeq. Similar yields of raw, mapped, filtered and deduplicated reads were obtained by both methods. F) Percentage of useful bins per library. Mapped, filtered and deduplicated reads were counted in 5 Kb non-overlapping windows and percentages of windows with and without reads were calculated. Dotted vertical lines define the threshold to define high-quality sequenced libraries (≤20 of empty windows).

### Optimized library preparation for Repli-seq analysis

DNA fragmentation and library preparation for RT analysis are critical steps to produce high-quality sequencing data. However, sample loss is also observed during these steps due to multiple sample transfers between DNA fragmentation, DNA clean-up, end-repair, adapter ligation and indexing. In addition, current Repli-seq methods rely on DNA sonication to produce fragments for efficient library preparation and sequencing. Ultrasonicators, such as Covaris or Bioruptor, are commonly used due to their consistency in producing tight fragment size distributions from a broad range of DNA concentrations (Marchal et al. 2018; Hayakawa et al. 2021). However, these methods require expensive consumables and equipment typically housed in core facilities. Here, we incorporated a single-tube DNA fragmentation and library preparation that reduce sample transfers, DNA loss and does not require expensive equipment and consumables (**Fig. 2A**). Distinct library preparation kits that incorporate DNA enzymatic fragmentation are currently available and all should produce comparable results. However, we optimized our protocol using the KAPA HyperPlus library prep kit, due to its high conversion rates of input DNA to adaptor-ligated fragments. We compared DNA libraries produced by the standard and optimized methods to validate our optimized protocol. First, we compared fragment distribution of libraries produced by enzymatic fragmentation with those produced by the standard Covaris sonication. Distinct incubation times were tested, and we found that 30-40 minutes of enzymatic fragmentation produced similar fragment size distributions as those obtained with Covaris sonicator (**Fig. 2D**). Next, end-repair and adapter ligation were performed according to the manufacturer instructions and DNA was purified and immunoprecipitated with an anti-BrdU antibody. Then, we sequenced the libraries produced by standard and optimized methods and compared the quality of sequencing data. We found similar yields of raw, mapped, quality filtered and final number of reads after removing PCR duplicates, with >83 % reads on average remaining for all samples sequenced (**Fig. 2E**). In this analysis, libraries were sequenced in the Illumina NovaSeq using 50bp of pair-end sequencing; however, we did not observe differences in mappability between single-end or pair-end sequencing. Binning sequencing reads into distinct size windows is required to calculate RT. Thus, we counted reads per fraction in 5 Kb non-overlapping windows (standard binning size used in Repli-seq). High quality sequencing data is expected to produce useful windows (bins containing sequencing reads) covering at least 80 % of the genome (windows with no reads are expected for highly repetitive sequences, such as centromeric and telomeric regions). Comparison of the fraction of windows with reads showed comparable data quality obtained from the standard and optimized protocols, with windows containing reads representing >90% of the reference genome (**Fig. 2F**). These results confirmed that single-tube DNA fragmentation and library preparation produced high-quality sequencing data.

### Optimized Repli-seq produces high quality RT data

Once we confirmed increased yields of input DNA, as well as high-quality of sequencing data using our optimized protocol, we generated RT profiles from H9, U2OS and MEFs cell lines (**Fig. 3**). First, we compared the raw RT data, i.e., unprocessed Log2(E/L) values, to evaluate the quality of the data produced by both methods (**Fig. 3A**). To do so, samples were processed using the optimal conditions tested and validated using the standard Repli-seq protocol: 2 hours of BrdU labeling, FACS sorting of 20,000 cells in each early and late S-phase fractions, tight fragment distribution, limited cycles of library amplification (10-11), and sequencing at ~8M reads per sample library. We found high-quality RT data from both methods for all cell lines analyzed (**Fig. 3A**). The best approach for quality assessment of RT data is to calculate the autocorrelation function (ACF), which is the correlation between the closest neighbor data points in each dataset. Since neighboring chromosomal regions are expected to replicate at the same time, the ACF values of high-quality datasets should be high. High quality datasets typically have ACF values > 0.6 (Ryba et al. 2011; Marchal et al. 2018). However, we have successfully characterized the RT program from patient samples with ACF values as low as 0.4 (Sasaki et al. 2017; Rivera-Mulia et al. 2019b). Here, we obtained very high-quality datasets using both methods, with ACF values > 0.9 and a slight improvement using our optimized method (**Fig. 3B**). To be able to compare RT datasets, a common practice is to quantile-normalize and smooth (local polynomial smoothing – LOESS) the data to remove noise and re-scale distinct datasets to equivalent dynamic range. Thus, we scaled and smoothed the RT datasets produced by the two methods. We found identical RT profiles for all samples analyzed regardless of the method used (**Fig. 3C**). Previously, we established that significant differences in RT between samples are > 1.5 (for scaled datasets with dynamic range of −6 to 6), while differences between technical and biological replicates are < 1.5 (Rivera-Mulia et al. 2015, 2017, 2019b). Here, we measured the differences in RT values across the genome between RT datasets produced by the standard or optimized methods. We found no significant differences when comparing RT datasets from the same cell type (**Fig. 3D**). As control, we compared RT datasets from distinct cell types and found significant differences in >20% of the genome (**Fig. 3D**). Finally, we performed a genome-wide correlation analysis of the RT datasets produced by the standard and optimized Repli-seq methods. We found correlation ≥0.99 between datasets from the same cell type regardless of method used, while correlations between different cell types were ≤0.77 (**Fig. 3E**). Altogether, our results confirm that our optimized Repli-seq method produces high quality RT data that is directly comparable with that obtained using the standard method.

**Fig. 3.**
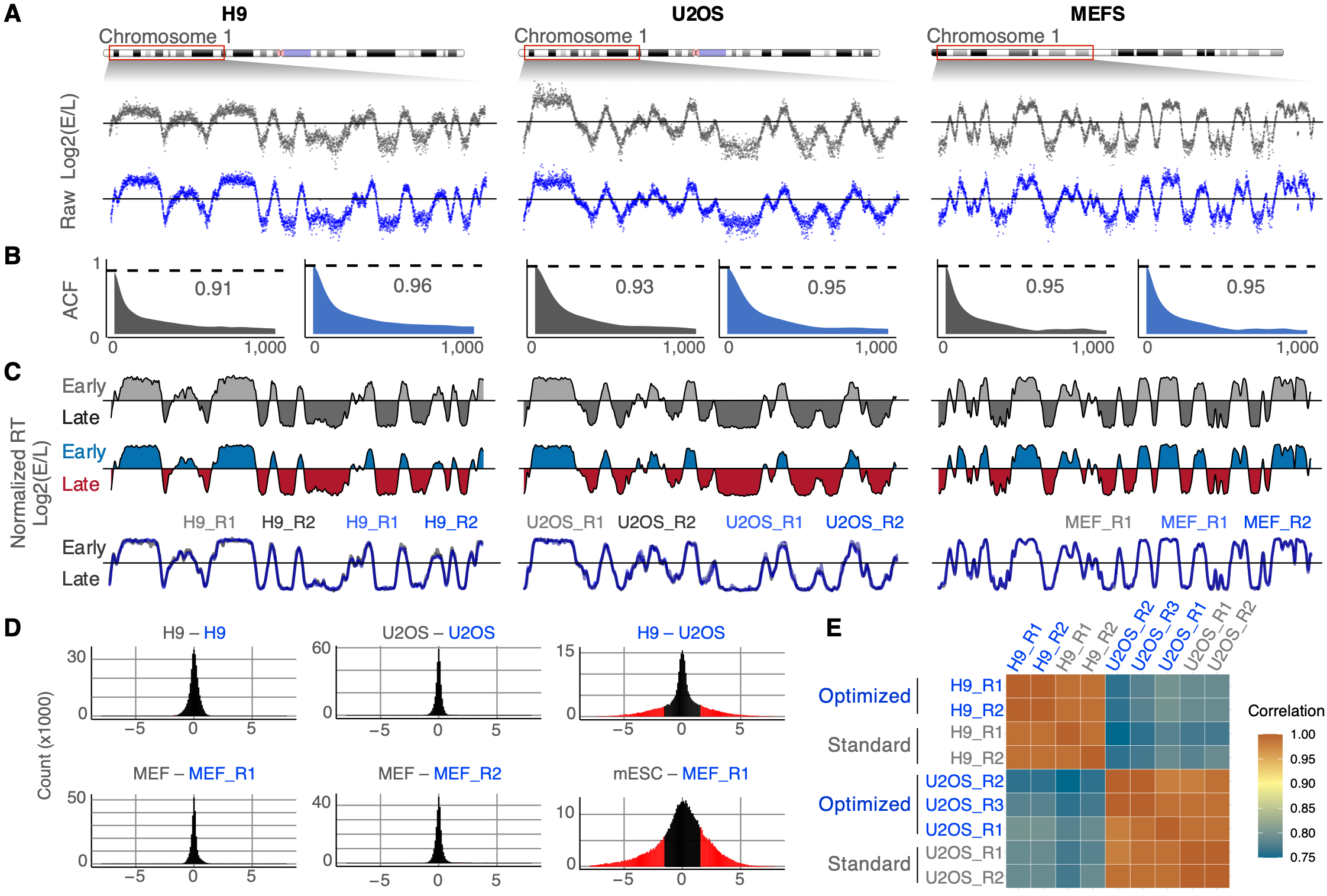
High-quality RT data is obtained by the optimized Repli-seq method. A) Raw RT data obtained from three distinct cell lines by the standard and optimized Repli-seq methods. 20K cells from each early and late S-phase fractions were processed. Raw RT data (unprocessed Log2 E/L ratio) is shown for a chromosomal region in H9 hESCs, U2OS cells and MEFs. B) Quality assessment of RT data. High quality data was obtained by both methods according to the ACF values (>0.9). ACF plots from 0-1,000 lag are shown. Dotted line shows the ACF at lag2 (closest neighbor data point) and ACF value is shown. C) Identical profiles are produced by the standard and optimized Repli-seq methods. Standard RT profiles are shown in grey and optimized RT profiles are shown in blue-red. Overlay plots show the high similarity of RT profiles obtained by both methods. Two replicates of each method are shown. D) Analysis of RT differences. No significant differences were observed between standard and optimized Repli-seq methods. Subtraction of RT data between standard and optimized methods were performed, as well as differences between distinct cell types. RT differences ≥1.5 are shown as red bars in the histograms. E) Pearson’s correlation analysis between standard and optimized RT profiles. Genome-wide correlation of RT programs was calculated for all samples and replicates processed.

### Optimal window size determination for Repli-seq analysis

We found that the RT data produced by our optimized Repli-seq protocol is comparable to that obtained with the conventional method using the standard data processing parameters. However, data processing parameters are selected arbitrarily and no systematic analyses to determine optimal parameters have been performed. In fact, there is no consensus on the binning size used to process RT datasets, with window sizes ranging from 5 kb to 1Mb (Marchal et al. 2018; Rivera-Mulia et al. 2018a; Massey et al. 2019; Rivera-Mulia et al. 2019b). Here, we analyzed the RT data produced with distinct binning sizes to determine the optimal windows to collect the reads. We counted reads in windows of 5, 10, 20, 50 and 100 kb and calculated the fraction of empty windows. We did not find significant differences using distinct window sizes when processing samples with sufficient number of cells (>10,000 cells per fraction), and consistently obtained >90 % of windows with RT values (**Supplementary Figure 1**). However, when processing limited samples (≤1,000 cells per fraction), up to 40 % of the smallest windows were lost as they did not contain reads (**Supplementary Figure 1**). Thus, to determine the optimum window size, we calculated the ACF values obtained from datasets with different window sizes. ACF values were consistently >0.9 for samples with >20,000 cells, however they were significantly improved with increasing window sizes for datasets obtained with limited cell numbers (**Supplementary Figure 1**). Since no significant decrease in empty windows was observed for binning the data in windows >20 kb, and ACF values were high at this binning size for datasets with distinct cell numbers, we established 20 kb as the optimal window size for processing sequencing reads for RT analysis. In addition, replication domains range in size from 400 kb to 1.2 Mb (Hiratani et al. 2010; Rivera-Mulia et al. 2015). Thus, a window size of 20 kb allows the generation of high-quality data without losing biological variation in DNA replication by averaging data points from different replication domains.

### Optimal smoothing parameters determination for Repli-seq analysis

Another important parameter for RT data analysis is the degree of smoothing to remove noise. LOESS smoothing is broadly used with distinct spans adjusted proportionally to the chromosome lengths (Hiratani et al. 2008; Ryba et al. 2011; Marchal et al. 2018; Rivera-Mulia et al. 2019b; Hadjadj et al. 2020; Koren et al. 2021). However, this parameter is also selected arbitrarily and usually adjusted after visual inspection of the RT profiles obtained. Here, we used the ACF calculation to determine the optimal LOESS span parameter to obtain comparable RT datasets. To do so, we generated RT datasets from samples with distinct numbers of cells at distinct degrees of smoothing (LOESS span calculated from 100 kb to 1.25 Mb divided by the chromosome length). For samples with ≥ 20,000 cells, we found highly similar RT profiles regardless of the degree of smoothing (**Supplementary Figure 2A**). However, we found a significant improvement increasing the smoothing for datasets generated from samples with limited cell numbers (**Supplementary Figure 2A**). LOESS smoothing produces neighbor data points with highly similar values; then, ACF values are usually very high after smoothing RT data (ACF at lag2 ≥ 0.99). Thus, to avoid over-smoothing that would cause loss of biological variation in the data, we calculated the ACF at distinct lags. Specifically, we calculated the ACF at lags corresponding to 100 and 200 kb. We found higher ACF values with increasing smoothing for all datasets (**Supplementary Figure 2B**). We defined the optimal smoothing parameter as the point when no further increase in ACF values were observed (**Supplementary Figure 2B**). The establishment of the optimal smoothing degree allowed us to generate comparable RT datasets from samples with distinct cell numbers (see below).

### Optimized Repli-seq allowed RT analysis from low input samples

The standard Repli-seq method is very robust and allows the generation of highly reproducible data. In fact, correlations between technical replicates are expected to be ≥0.95 and correlations between biological replicates, even from distinct individuals, are ≥0.90 for high quality datasets (Rivera-Mulia et al. 2017; Marchal et al. 2018; Rivera-Mulia et al. 2019b). However, reproducibility of the RT measurements strongly depends on the availability of thousands of proliferating cells. The standard Repli-seq method is performed with 20,000 to 80,000 cells per S-phase fraction. Although we have obtained useful RT data with as low as 1,000 cells per fraction, the quality of the RT profiles decreased significantly (ACF values ≥0.4) and stronger smoothing was required to reduce noise (Sasaki et al. 2017; Marchal et al. 2018; Rivera-Mulia et al. 2019b). Thus, here we tested the capacity of our optimized method to analyze RT from very low input cells. To do so, we sorted distinct numbers of cells ranging from 40,000 cells and as low as 100 cells into early and late S-phase fractions. RT profiles from these samples were generated in duplicates. Decreased yields of purified DNA were observed with decreasing number of cells, and although DNA was outside of the detection limits both by photometric (Nanodrop) and fluorometric (Qubit) measurements, samples were processed, and libraries constructed (**Supplementary Figure 3A**). To ensure successful conversion of input fragments to adapter-ligated libraries, adapters were diluted as shown in the **Supplementary Table 1**. To produce sufficient amounts of DNA libraries, the number of PCR cycles was optimized according to the DNA input (**Supplementary Table 2**). Repli-seq libraries were sequenced at ~8M reads per library and 50bp of pair-end sequencing in the Illumina NovaSeq. We found high yields of mapped, filtered and deduplicated reads for libraries derived from 5,000 to 40,000 cells (**Supplementary Figure 3B**). However, libraries produced from ≤1,000 cells per fraction resulted in high rates of PCR duplicates (average of 35% reads remaining after PCR duplicate removal), due to the high number of PCR cycles required to generate enough material for sequencing. Nevertheless, we found that the remaining fraction of mapped/deduplicated reads was sufficient to assess the RT. First, we analyzed the quality of raw RT data, obtained by calculating the Log2(E/L) ratio of reads binned into non-overlapping 20 kb windows. We found very high-quality RT datasets with as low as 500 cells (ACFs >0.8) and good RT datasets with only 100 cells per fraction (ACF ≥ 0.58), confirming that our optimized method can be applied for very low input samples (**Fig. 4A-B**). Next, we compared the normalized and smoothed datasets using the optimal parameters described above. We found highly similar results for RT datasets obtained from standard and very low input samples (**Fig. 4C**). In fact, overlay plots show identical RT profiles at distinct chromosomal regions for all samples generated from 40,000 cells or as low as 100 cells (**Fig. 4D**). Moreover, correlation analysis demonstrated the high similarity between RT datasets produced with different numbers of cells ranging from 40,000 cells and as low as 100 cells (**Fig. 4E**). Overall, our results validate our optimized protocol and demonstrate its applicability for assessing the RT program of very limited samples without compromising the quality and resolution of the data produced.

**Fig. 4.**
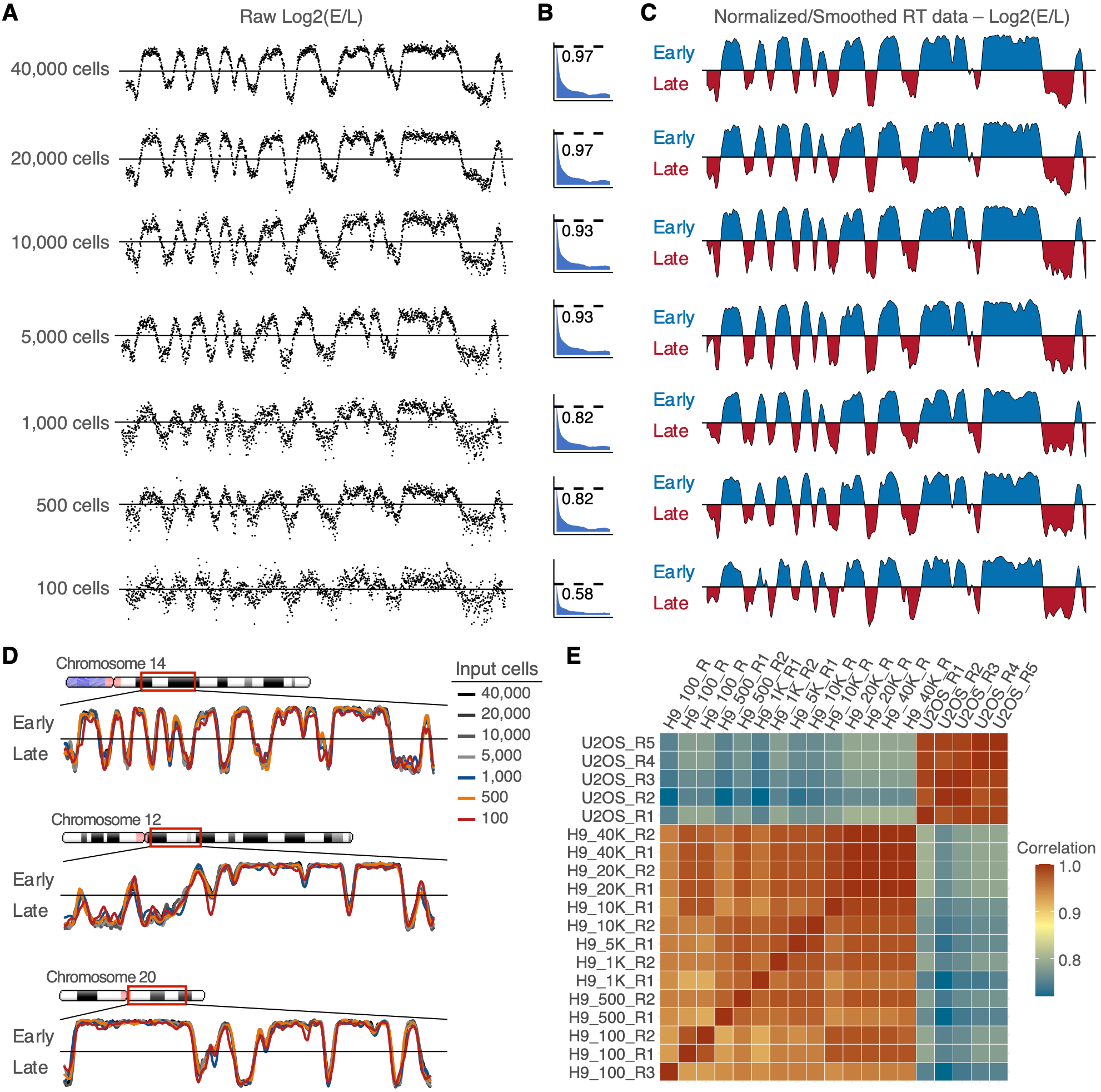
Optimized Repli-seq allows RT profiling from low input samples. A) Raw RT data (Log2 E/L values) were obtained from 40K, 20K, 10K, 5K, 1K, 500 and 100 cells. RT values were calculated at non-overlapping 20Kb windows. B) Quality assessment of RT data obtained from standard (≥20K cells) and low input samples (≤10Kb). ACF plots from 0-1,000 lag are shown. Dotted line shows the ACF at lag2. C) High similarity of normalized RT profiles was obtained from standard and low input samples. RT data was quantile normalized and LOESS smoothed as described in the methods. D). Identical, high quality RT profiles were obtained from standard and low input samples. RT profiles from 40K, 20K, 10K, 5K, 1K, 500 and 100 cells are shown for distinct chromosomal regions. E) Correlation analysis of RT programs derived from two human cell lines obtained from standard and low input samples. Correlations ≥0.93 were obtained from input samples ranging from 100 to 40,000 cells per fraction. The U2OS cell line was used as a control of a distinct cell type (correlations between U2OS and H9 RT datasets were ≤0.77).

## Conclusion

In this paper, we present an optimized Repli-seq method that allows to profile replication timing genome-wide. Current protocols are robust and highly reproducible but restricted to samples with thousands of proliferating cells, are highly laborious, time consuming and expensive. Moreover, the multiple tube-to-tube transfers throughout the protocol, from cells to the purified DNA, lead to a significant sample loss. Thus, we developed an improved method that reduces sample loss, results in increased DNA yields, and provides high quality DNA libraries for analysis of RT by next-generation sequencing. Our optimized Repli-seq method minimizes the number of sample transfers by sorting cells directly into the lysis buffer, removing the digestion step with proteinase K, and performing a single-tube DNA fragmentation and library preparation. The increased nucleic acid yields obtained with our optimized Repli-seq protocol allows us to obtain high quality RT data from very low input samples (as low as 100 cells per fraction of early and late S-phase populations). Our improved method also allows rapid purification of DNA from proliferating cells, reducing the processing time from hours to minutes. In addition, we provided a standardized bioinformatic pipeline to process RT data obtained from distinct numbers of cells. Optimization of data processing allowed generation of high-quality data of low input samples that could be directly compared against standard input samples. Finally, the optimized Repli-seq protocol exploits enzymatic fragmentation, removing the need of expensive equipment and consumables, making the measurement of RT accessible to a broader range of researchers.

## Material and methods

### Cell lines

H9 cells were cultured in mTeSR Plus (StemCell Technologies Cat. No. 05825) on Geltrex (Life Technologies Cat. No. A1413302) coated dishes according to WiCell instructions. Cells were dissociated with Accutase (Life Technologies Cat. No. A011105-01) for expansion. U2OS cells were grown in DMEM media (Corning Cat.No. 15013CM) supplemented with 10% FBS (Corning Cat.No.35011CV), 1% Glutamine (Gibco Cat.No. 25030081), 1% Penicillin-Streptomycin (Gibco Cat.No. 15070-063). MEFs were cultured in 1:1 DMEM media (Corning Cat.No. 10-017-CV) and F10 media with Glutamax (Gibco Cat.No. 41550021), supplemented with 10% FBS (Sigma Cat.No. F2442), 1X NEAA (Gibco Cat.No. 11140050) and 1% Penicillin-Streptomycin (Corning Cat.No. 30002Cl). At ≤70% confluency cells were labeled and fixed for Repli-seq as previously described (Ryba et al. 2011; Marchal et al. 2018). Briefly, cells were pulse-labeled with 100 uM BrdU for 2 hours. Single-cell suspensions of BrdU-labeled cells were harvested, fixed in 70% ethanol and stored at −20 °C.

### Flow cytometry

2 x 10^6 BrdU-labeled and fixed samples were resuspended in PBS with 1% (vol/vol) FBS, stained with 50ug/ml PI and treated with 250ug/ml RNase-A, filtered with a 36 um nylon mesh, and FACS sorted into 1.5 mL tubes. FACS sorting of specific cell numbers was performed on a FACSAria at the University Flow Cytometry Resource or in our laboratory SONY SH800 cell sorter. For the standard Repli-seq, sorted cells were collected into 400 uL of PBS/FBS and mixed thoroughly. For the optimized Repli-seq, sorted cells were collected into 400 uL of the lysis buffer from the Zymo Quick-DNA purification kit (Cat. D3021) and mixed thoroughly.

### DNA purification from sorted cells

DNA was isolated using the Zymo Quick-DNA purification kit (Cat. D3021). For the standard Repli-seq, sorted cells were centrifuged at 400g, resuspended in SDS buffer (2.5 ml of 1 M Tris-HCl, pH 8.0, 1ml of 0.5 M EDTA, 10 ml of 5 M NaCl and 2.5 ml of 10% SDS), digested 2 hours with 0.2 mg/ml proteinase K, 800 ul of genomic lysis buffer were then added from the Zymo Quick-DNA kit and DNA was purified according to the manufacturer instructions. For the optimized Repli-seq, sorted cells were collected directly into 400 uL of the lysis buffer from the Zymo Quick-DNA kit and DNA was purified according to the manufacturer instructions. For the optimized Repli-seq samples can be stored in the genomic lysis buffer at −20 °C for months.

### Standard Repli-seq library preparation

Purified DNA from sorted cells was fragmented in the Covaris sonicator as previously described (Marchal et al. 2018). Briefly, purified DNA was transferred to microtube AFA fiber and sonicated with 175 W, 10% duty cycle, 200 cycles per burst, 120 s at 4 °C. Fragmented DNA was then concentrated using the Zymo DNA Clean & Concentrator-5 kit (Cat. D4014). Then libraries were constructed using the NEBNext Ultra DNA Library Prep Kit for Illumina (NEB, Cat. No. E7370) according to the manufacturer instructions. Libraries were concentrated using the Zymo DNA Clean & Concentrator-5 kit. BrdU-IP was performed after library construction using 2.5 μg/ml anti-BrdU antibody (BD, cat. 555627) as previously described (Marchal et al. 2018). Immunoprecipitated libraries were indexed and amplified using multiplex oligos from NEB. Amplified libraries were purified using AMPure XP beads (Cat. No. XXX) according to the manufacturer instructions and quality controls were performed before sequencing (Qubit quantification of DNA concentration, size distribution estimation on Agilent Bioanalyzer or TapeStation, and enrichment validation of known markers by PCR-Ryba et al, 2011).

### Optimized Repli-seq library preparation

Purified DNA from sorted cells was directly processed using the KAPA HyperPlus library preparation kit (Roche Cat. No. 07958978001) according to the manufacturer instructions. This method allows a single-tube DNA fragmentation and library preparation. Enzymatic fragmentation was performed at 37 °C for 30 minutes, followed by end-repair and ligation of KAPA dual-indexed adapters (Roche Cat. No. 08278555702). Libraries were purified using the Zymo DNA Clean & Concentrator-5 kit and processed for BrdU-IP as described above. Amplification of immunoprecipitated libraries was performed using the KAPA HiFi HotStart from the KAPA HyperPlus kit according to the manufacturer instructions. Amplified libraries were purified using AMPure XP beads according to the manufacturer instructions and quality controls were performed before sequencing as described above.

### Next-generation sequencing

Libraries were pooled at a final concentration of 10 nM and DNA concentration and size distribution from each pool were confirmed by Qubit quantification and size estimation on Agilent Bioanalyzer or TapeStation. Library pools were sequenced at University of Minnesota Genomics Center (UMGC), on the Illumina NovaSeq at ~8M reads per library at 50 bp of pair-end sequencing.

### Data analysis and visualization

Data analysis was performed at the Minnesota Supercomputing Institute (MSI). Sequencing reads were trimmed for adapter sequenced using Cutadapt (Martin 2011) and aligned to the reference genome with Burrows-Wheeler Aligner (Li and Durbin 2010). Sequence Alignment Map (SAM) files were converted into Binary Alignment Maps (BAM) with SAMtools (Danecek et al. 2021). Quality filter and removal of PCR duplicates were performed with SAMtools using the samtools view and markdup functions respectively. Mapped, filtered and deduplicated reads were counted into genomic windows of distinct size (from 5 to 100 kb) using the intersect function from BEDtools (Quinlan and Hall 2010). RT data values were calculated as the Log2(E/L) ratios. Downstream processing, including quantile normalization and LOESS smoothing, and data visualization was performed in R (RCoreTeam 2017).

## Acknowledgements

We thank Laura Niedernhofer and Hai Dang Nguyen for providing the MEFs and U2OS cell lines; and Silvia Meyer-Nava and Anala Shetty for critically reading the manuscript.

## Author Contributions

JCRM conceived and designed the study. JCRM, CTG and SMC conducted experiments. JCRM analyzed data and wrote the manuscript. All authors read and approved the manuscript.

## Data Availability

Datasets generated in this study are available in the Gene Expression Omnibus (http://www.ncbi.nlm.nih.gov/geo/), under the number GSE196749.

