## Supplementary Material for "Optimized Repli-seq: Improved DNA Replication Timing Analysis by Next-Generation Sequencing"

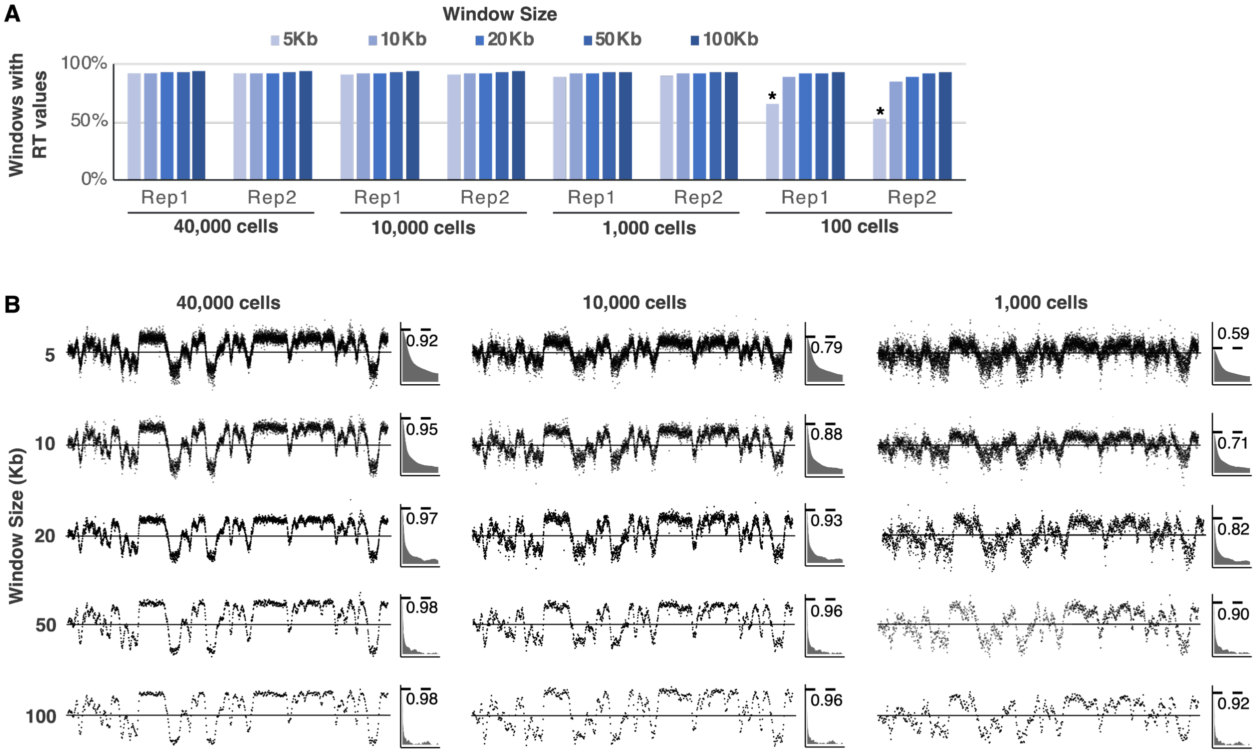

**Supplementary Figure 1**

Optimal window size determination for Repli-seq analysis. A) Windows with RT values were quantified for RT datasets derived from distinct cell inputs (40,000, 10,000, 1,000 and 100 cells), and produced with distinct binning sizes (5, 10, 20, 50 and 100 kb). Two replicates of each sample are shown. B) Analysis of RT at distinct window sizes. Raw RT Log2(E/L) values are shown for datasets derived from 40,000, 10,000, and 1,000 cells. Windows sizes from 5 kb to 100 kb were tested. ACF plots from 0-1,000 lag are shown. Dotted line shows the ACF at lag2 (closest neighbor data point) and ACF value is shown.

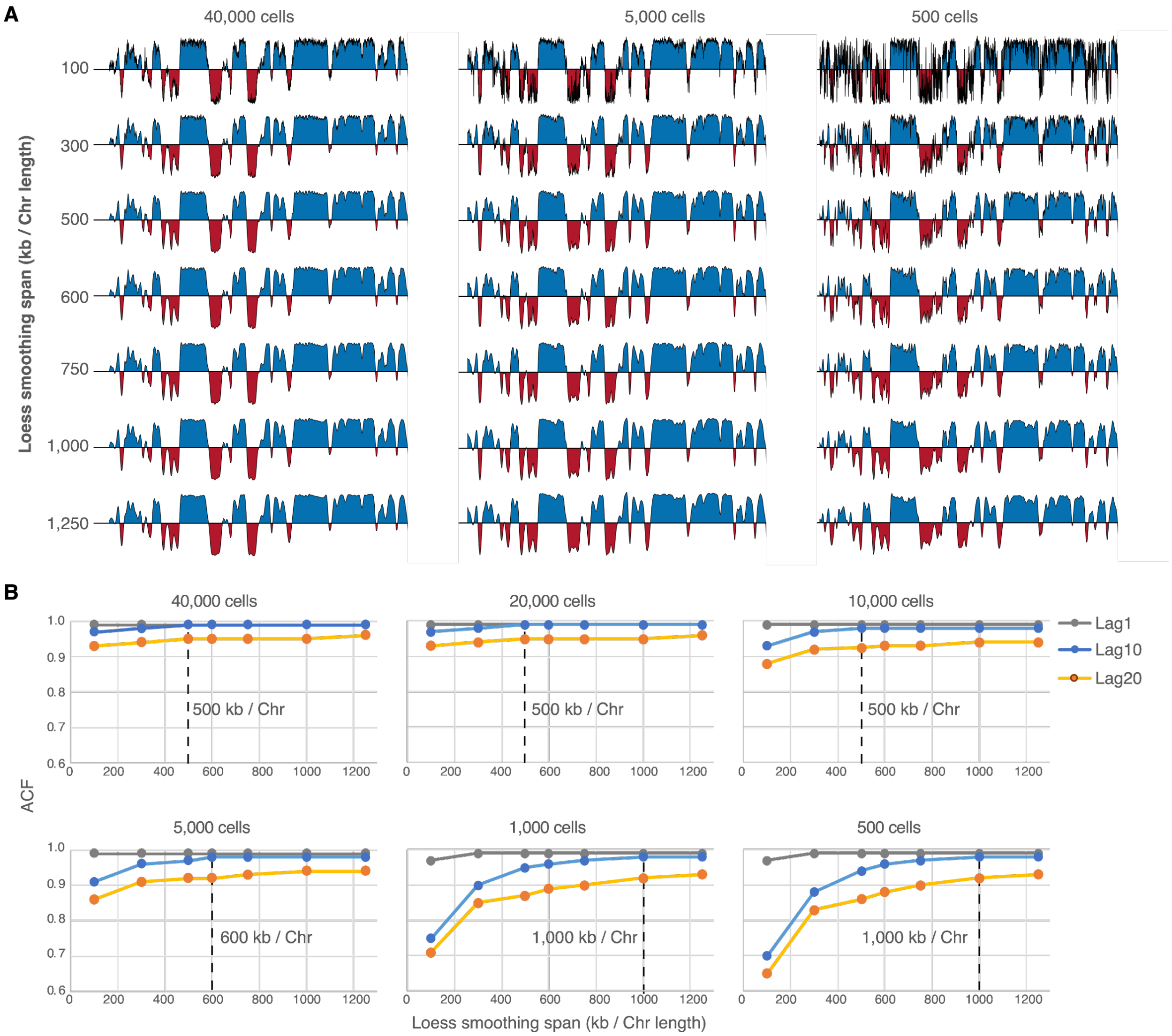

**Supplementary Figure 2**

Optimal smoothing determination for Repli-seq analysis. A) RT datasets derived from samples with distinct numbers of cells were generated. Distinct degrees of smoothing were tested (LOESS span calculated from 100 kb to 1.25 Mb divided by the chromosome length). B) ACF values at distinct lags (1-20) were calculated for RT datasets generated with span from 100 kb to 1.25 Mb. Dashed vertical line indicates the optimal span for LOESS smoothing for each dataset.

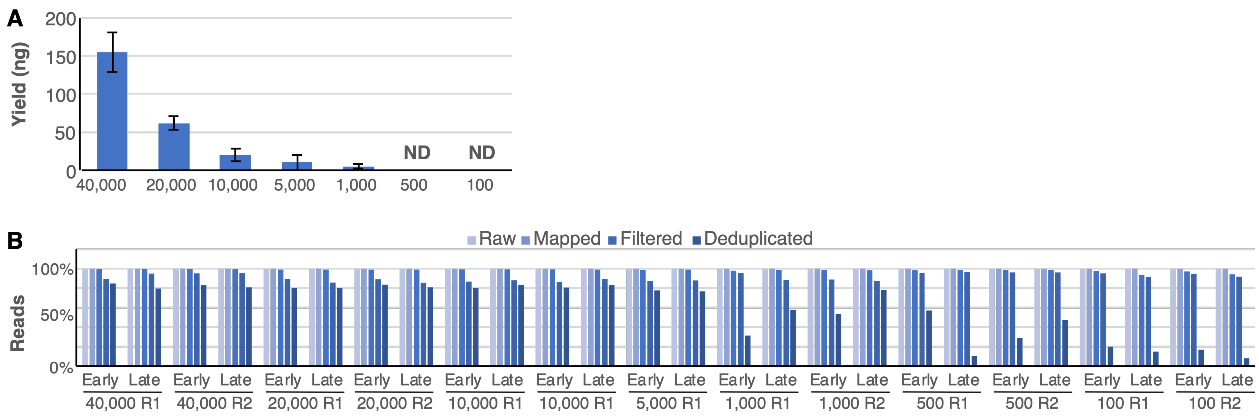

**Supplementary Figure 3**

A) Genomic DNA yields of distinct number of sorted cells. Bars represent the average of 4 replicates and standard deviation is shown. B) Comparison of NGS reads obtained from distinct input cells by the optimized Repli-seq method. Distinct number of cells, ranging from 40,000 to 100 cells, of early and late S-phase cells were collected by flow cytometry. All samples were sequenced at ~8M reads per library using pair-end sequencing at 50 bp in the Illumina NovaSeq. Yields of raw, mapped, filtered and deduplicated reads are shown.

| Input DNA | Adapter [ ] | Adapter:Fragment Ratio | Dilution | Notes |
|---|---|---|---|---|
| 100-200 ng | 15 μM | ~40:1 | NA | Range of standard yield of Repli-seq samples with ≥ 20K cells |
| 50-100 ng | 15 μM | ~100:1 | NA | |
| 5-50 ng | 5 μM | ~300:1 | 1:3 | Expected yields from 5K-10K cells |
| ≤ 5 ng | 1.5 μM | ~900:1 | 1:10 | Expected yields from < 5K cells |

**Supplementary Table 1**

Adapter dilution for optimized Repli-seq library preparation. To ensure high conversion rates of input fragments to adapter-ligated libraries, adapter:fragment ratios were adjusted according to the input DNA. For inputs ≥ 50 ng an adapter:insert molar ratio of between 40:1 to100:1 yields an optimum conversion efficiency. For low inputs (<50ng) an adapter:insert molar ratios between 300:1 to 1,000:1 yields better conversion efficiencies. Adding an increased amount of adapters in this step generates adapter-dimers. However, dimers and concatenates are removed during the BrdU-IP step, thus high adapter concentrations were used to enhance ligation efficiency.

| Input DNA before BrdU-IP | Number of cycles |
|---|---|
| 200 ng | 9-10 |
| 100 ng | 10-11 |
| 50 ng | 12-13 |
| 10 ng | 16-17 |
| 5 ng | 17-18 |
| < 5 ng | 18-19 |

**Supplementary Table 2**

Number of PCR cycles for Repli-seq library amplification. Repli-seq libraries were amplified using the KAPA HyperPlus kit and KAPA Library Amplification Primer Mix. To produce enough library DNA the number of cycles was adjusted according to the initial amount of input DNA. For samples derived from ≤ 1,000 cells DNA was outside of the detection limits and 19 cycles were used to amplify the libraries for these samples.
